# Detecting genetic effects on phenotype variability to capture gene-by-environment interactions: a systematic method comparison

**DOI:** 10.1101/2023.06.09.544298

**Authors:** Xiaopu Zhang, Jordana T. Bell

**Affiliations:** Department of Twin Research and Genetic Epidemiology, King’s College London, London, UK

## Abstract

Phenotypic variability has been widely observed across organisms and traits, including in humans. Both gene-gene and gene-environment interactions can lead to an increase in phenotypic variability. Therefore, detecting the underlying genetic variants, or variance Quantitative Trait Loci (vQTLs), can provide novel insights into complex traits. Established approaches to detect vQTLs apply different methodologies from variance-only approaches to mean-variance joint tests, but a comprehensive comparison of these methods is lacking. Here, we review available methods to detect vQTLs in humans, carry out a simulation study to assess their performance under different biological scenarios of gene-environment interactions, and apply the optimal approaches for vQTL identification to gene expression data. We find that the squared residual value linear model (SVLM) method is optimal when the interacting exposure is discrete, and both SVLM and the deviation regression model (DRM) perform well when the interacting exposure is continuous. Additionally, a larger sample size, smaller minor allele frequency, and more balanced sample distribution in different exposure categories increase power of SVLM and DRM. Our results highlight vQTL detection methods that perform optimally under realistic simulation settings and show that their relative performance depends on the type of exposure in the interaction model underlying the vQTL.

**Author Summary:** Genetic background can influence organismal response to changes in the environment, including through effects of gene-environment (GxE) interactions. GxE interactions form a fundamental component of complex phenotypes and disease. The presence of GxE interactions leads to increases in phenotypic variability, where individuals of a certain genotype can display more divergent phenotypes. Therefore, identifying genetic variation that can alter phenotypic variance is an alternative approach to identifying genetic variants that underlie GxE effects. Here, we conduct a systematic assessment of methods that detect genetic variants linked to phenotypic variance under several GxE scenarios. We identify the most optimal approaches for different environmental exposure settings and apply these to validate previously detected GxE signals. We also estimate power of these methods to detect GxE effects. Our results help guide experimental design for future studies aiming to identify genetic variant impacts on phenotypic variance, and their result interpretation in context of gene-environment interactions in complex traits.

## Introduction

Phenotypic variability has been observed across a broad range of organisms and phenotypes, including for example in human obesity[1,2]. One driver of phenotypic variability is gene-gene (GxG) and gene-environment (GxE) interactions. These represent situations in which individuals of a certain genotype can display more divergent phenotypes due to genotype interactions with specific exposures or genetic contexts[2,3,12,4–11]. Previous studies have proposed that GxG and GxE interactions commonly influence complex traits including human diseases[13–16], animal and plant breeding outcomes[17–20], and evolutionary traits[21–29]. For example, a GxG interaction between genetic variants in *ERAP1* and *HLA-C* is implicated in susceptibility to psoriasis[16], and a smoking-genotype interaction in *NAT2* is associated with bladder cancer risk[30].

Genetic variants associated with phenotypic variability are defined as variable quantitative trait loci, or vQTLs (Fig 1A). Genetic variants involved in GxG or GxE interactions are one type of vQTLs. Therefore, detecting vQTLs could uncover regulatory mechanisms underlying complex traits. Compared to directly identifying GxG and GxE effects, vQTLs identification is more expedient because it does not require a comprehensive multi-dimensional search for interactions among multiple factors, or previous knowledge about potential interactions between specific exposures and targeted genetic variants[7,20,31].

**Fig 1.**
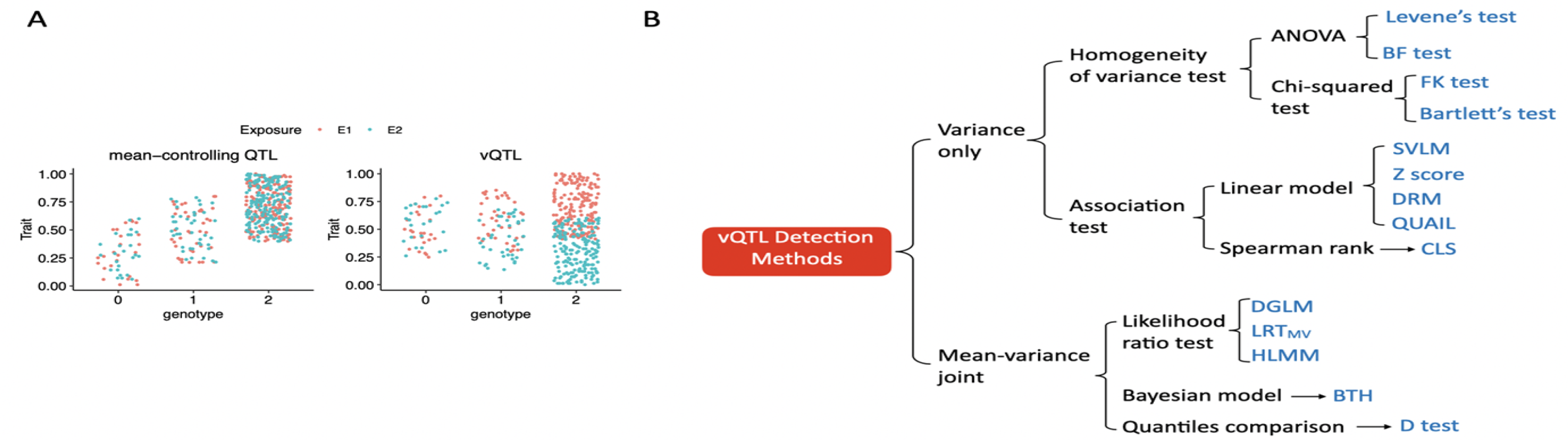
vQTL detection method overview. A) Example of traits associated with a mean-controlling QTL (left) and a vQTL (right). B) Overview of vQTL identification methods. Joint location and scale tests which require external annotations of detected QTLs are not included.

Multiple vQTL studies have been conducted in humans and across a wide range of quantitative and discrete traits (Table 1). Yang et al (2012) discovered a genetic variant in *FTO* to be significantly associated with BMI variability[1]. More recently, Wang et al. (2019) identified 75 vQTLs to influence phenotype variability for at least one of nine quantitative traits, including BMI, bone mineral density, and birth weight[2]. Several case-control studies have also identified vQTLs associated with human diseases[13–16], for example, rheumatoid arthritis[13]. In terms of intermediate phenotypes, vQTLs in humans have been reported for gene expression[32–34], DNA methylation[35,36], and protein level variability[31,37]. Further vQTL findings have also been obtained for multiple biochemical[38], physical[39,40], facial[41], and morphological traits[42].

**Table1:** Genome-wide vQTL studies in human samples.

| Trait [platform] | Number of samples | Genetic variants | Input data processing | vQTL detection<br>method | Significance<br>threshold | Findings |
| --- | --- | --- | --- | --- | --- | --- |
| **Intermediate traits** |  |  |  |  |  |  |
| DNA methylation profiles of 431,813 probes [Illumina 450K][35] | 729 unrelated blood samples | ~3.4 million SNPs within 500kb of CpG sites | Rank transformed (Bartlett's) and untransformed (Levene's test) data | Bartlett's and Levene's tests | Bonferroni correction (0.05): $P < 9.4e-11$ | 590 CpGs influenced by cis-vmeQTLs, and 14/590 CpGs were affected by GxE. 86% of vmeQTLs were still significant in Levene's test. |
| Gene expression levels of 35,078 probes [Illumina HT-12 V3][32] | 825 adipose, 825 LCL, and 705 skin samples from twins | SNPs within 1Mbp around each probe, ~1.25 million SNP-probe pairs | Coefficient of variance | FK test and DGLM | $P < 1e-8$ | Overall, 99, 75, and 59 genes influenced by v-eQTL in LCLs, fat, and skin respectively, and 8 genes are shared across tissues. |
| Gene expression levels of 13,660 genes [RNA-seq][33] | Discovery set: 765 LCL samples from twins<br>Replication set: 462 unrelated LCL samples | SNPs within 1Mbp around the TSS | Squared residuals after regressing out genotype effects | Spearman's rank correlation between residuals and genotype | FDR (0.05; 5 permutations), and Bonferroni correction (0.05): $P < 8.9e-10$ | FDR < 0.05 threshold: 497 v-eQTLs detected and 28/497 v-eQTLs replicated; 181 v-eQTLs are also eQTLs.<br>Bonferroni correction: 23 v-eQTLs detected, and 16/23 v-eQTLs replicated. |
| Gene expression levels of 15,574 genes [WG-6 version 1] and 12,076 genes [U133A array], where 10,188 genes were shared[60] | 210 LCL samples from 3 populations | SNPs within 1Mbp around the TSS; 747,524 and 763,745 SNP-probe pairs in the two sets | Coefficient of variance | DGLM and FK test | $P < 1e-8$ | 166 and 60 genes are associated with at least one v-eQTL respectively in the two datasets, and 8 genes are shared between the two datasets. |
| Gene expression levels of 9,957 protein-coding genes [single cell RNA-seq][34] | 5,447 iPSCs from 53 unrelated samples of African ancestry | 8.47 million SNPs within 100kb around the TSS | Variance, coefficient of variance, ratio of variance to mean, dispersion parameter | Linear model | Benjamini–Hochberg (0.1) | 235 eQTL and 5 v-eQTL detected. 85% of eQTLs are vQTLs; 100% vQTLs are eQTLs. |
| Protein levels of C-reactive protein and ICAM-1 [High-sensitivity CRP][31] | 21,799 unrelated blood samples | ~0.34 million SNPs | Log-transformed CRP levels | Levene’s test | $P < 1.5e-7$ | 1 vQTLs for C-reactive protein level variability, and 2 vQTLs for soluble ICAM-1 protein level variability. |
| **Phenotype traits** |  |  |  |  |  |  |
| 13 physical traits[2] | 348,501 unrelated samples | ~5.55 million SNPs | Adjusted and standardized phenotypes | Brown-Forsythe test | $P < 2e-9$ | 75 vQTLs identified for 9 traits; 66/75 vQTLs are mean-controlling QTLs. Altogether, 16/75 vQTLs show GxE. |
| BMI, height[1] | Discovery set: 133,154<br>unrelated samples<br>Replication set: 36,727<br>unrelated samples | ~2.68 million SNPs | Rank-based inverse-<br>normal transformed data | Linear model | $P < 5e-8$ | 6 vQTLs for height and 7 vQTLs for BMI; 1<br>vQTL for BMI is replicated. |
| BMI, height[39] | Discovery set: siblings<br>from 583 families,<br>Replication set:<br>monozygotic and dizygotic<br>twins from discovery set<br>families | ~0.26 million SNPs | Standard deviation | Linear model | $P < 1e-5$ | 4 vQTLs for height reach suggestive threshold<br>($P < 1e-5$), but none replicate.<br>2 vQTLs for BMI reach suggestive threshold<br>($P < 1e-5$), and 1 vQTL replicates |
| BMI[40] | 344,201 unrelated samples.<br>80% in discovery set, and<br>20% in replication set | ~6.7 million SNPs | Dispersion of samples to<br>the median of each group | DRM | $P < 5e-8$ | 26 vQTLs, 3/26 are mean-controlling QTLs |
| 20 facial traits[41] | Discovery set: 2,447<br>unrelated European-ancestry samples<br>Replication set: 5,128<br>unrelated Asian-ancestry<br>samples | ~3.1 million SNPs | Untransformed<br>phenotype values | Brown-Forsythe<br>test | $P < 5e-8$ | No signals surpass $P < 5e-8$, but 10 vQTLs reach<br>$P < 5e-7$ suggestive threshold. |
| Rheumatoid arthritis[13] | 19,108 unrelated European and US samples (3,323 cases and 15,785 controls) | 107,144 autosomal SNPs | Case-control design | Brown-Forsythe test | $P < 5 \times 10^{-8}$ | 2 vQTLs detected, and 2 other signals reach suggestive threshold ($P < 5 \times 10^{-7}$). |
| Psoriasis[14] | Discovery set: European samples, including 2,175 cases and 5,144 controls<br>Replication set: US samples, including 849 cases and 641 controls | Discovery set: ~5.5 million SNPs<br>Replication set: ~4.6 million SNPs | Case-control design | Brown-Forsythe test | $P < 5 \times 10^{-8}$ | 2 vQTLs detected in vGWAS, and GWAS. 5 other vQTLs reached suggestive threshold ($P < 2.5 \times 10^{-5}$) |
| 25 hydroxyvitamin D (25OHD) concentration[38] | 318,851 unrelated European-ancestry samples | ~6.09 million SNPs | Normalized adjusted 25OHD level | Brown-Forsythe test | $P < 5 \times 10^{-8}$ | 25 vQTLs detected, 23 of them are also mean-controlling QTLs while the remaining 2 are at GWAS suggestive threshold ($P < 10 \times 10^{-5}$) |
| 10 brain features[42] | 25,575 unrelated European-ancestry samples | >5 million SNPs | Rank-based inverse normal transformed data | Brown-Forsythe test | $P < 5 \times 10^{-8}$ | 1 vQTL detected for mean thickness and thalamus. |

\* LCL = lymphoblastoid cell line. FK = Fligner-Killeen test. DGLM = double generalized linear model. TSS = transcription start site. DRM =
deviation regression model.

A number of methods have been developed to identify vQTLs, and the general premise of these is to detect changes in phenotypic variance across different genotype groups. These methods can be categorized into variance-only and mean-variance joint tests, depending on whether they can detect both mean-controlling QTLs and vQTLs. Variance-only approaches include homogeneity of variance tests[43–47], a non-parametric approach[48], and association tests[1,33,40,49–51] (Fig 1B). Most mean-variance joint approaches use likelihood ratio tests to assess the role of genetic variants[52–57]. Further methods also include a Bayesian model[58] and a quantiles comparison[59] (Fig 1B). Several comparisons of published vQTL detection methods have been carried out, but these have been limited to up to 5 methods to date[2,51,58] or certain biological scenarios, for example, when the exposure is discrete[40].

Here, we carry out a simulation study to systematically compare the performance of ten vQTL mapping approaches under different biological scenarios. The simulation scenarios assume that different types of gene-environment interactions lead to phenotypic variability. We also evaluate the influence of data transformations on the methods’ performance, and we conduct a power analysis. Finally, we apply the optimal methods to validate previously detected vQTLs of human gene expression levels (v-eQTL).

## Results

We carried out a systematic comparison of ten previously proposed methods to detect variance QTLs. These include Levene’s test, BF test, FK test, Bartlett’s test, SVLM, DRM, Z score, CLS, QUAIL, and DGLM. Each method performance was evaluated using the false positive rate (FPR) and the discovery rate based on simulation results.

### vQTL detection methods comparison for discrete environmental exposures

We initially assumed that a GxE interaction leads to increased phenotypic variability and that the exposure is discrete, such as smoking (current smokers or not). We set a range of effect sizes of GxE interaction effects and errors were generated from Normal, Gamma, or Chi-squared distributions. The results are obtained for a sample size of 1000 unrelated individuals with a genetic variant at 0.05 MAF, and the sample proportions in the two exposure groups are 90% and 10%.

The Z score method shows high FPR across all error distributions (Fig 2A). The FPR of Bartlett’s test, CLS, DGLM, FK, Levene’s test, and QUAIL are near 0.05 when the error follows a Normal distribution, but greater when the error is from a non-Normal distribution (Fig 2A). The corresponding FPRs are higher when the error is from the Chi-squared distribution compared to errors from the Gamma distribution. These methods are sensitive to trait skewness because Chi-squared distributed errors lead to the greatest skewness of traits, but Normally distributed errors contribute the least to the trait skewness. The SVLM, BF, and DRM methods show a consistently low FPR whatever the error distribution (Fig 2A). These findings are consistent with previous comparisons which suggest that the FPR is higher for the DGLM and Z score regression methods, and lower for the SVLM and DRM methods[2,40].

**Fig 2.**
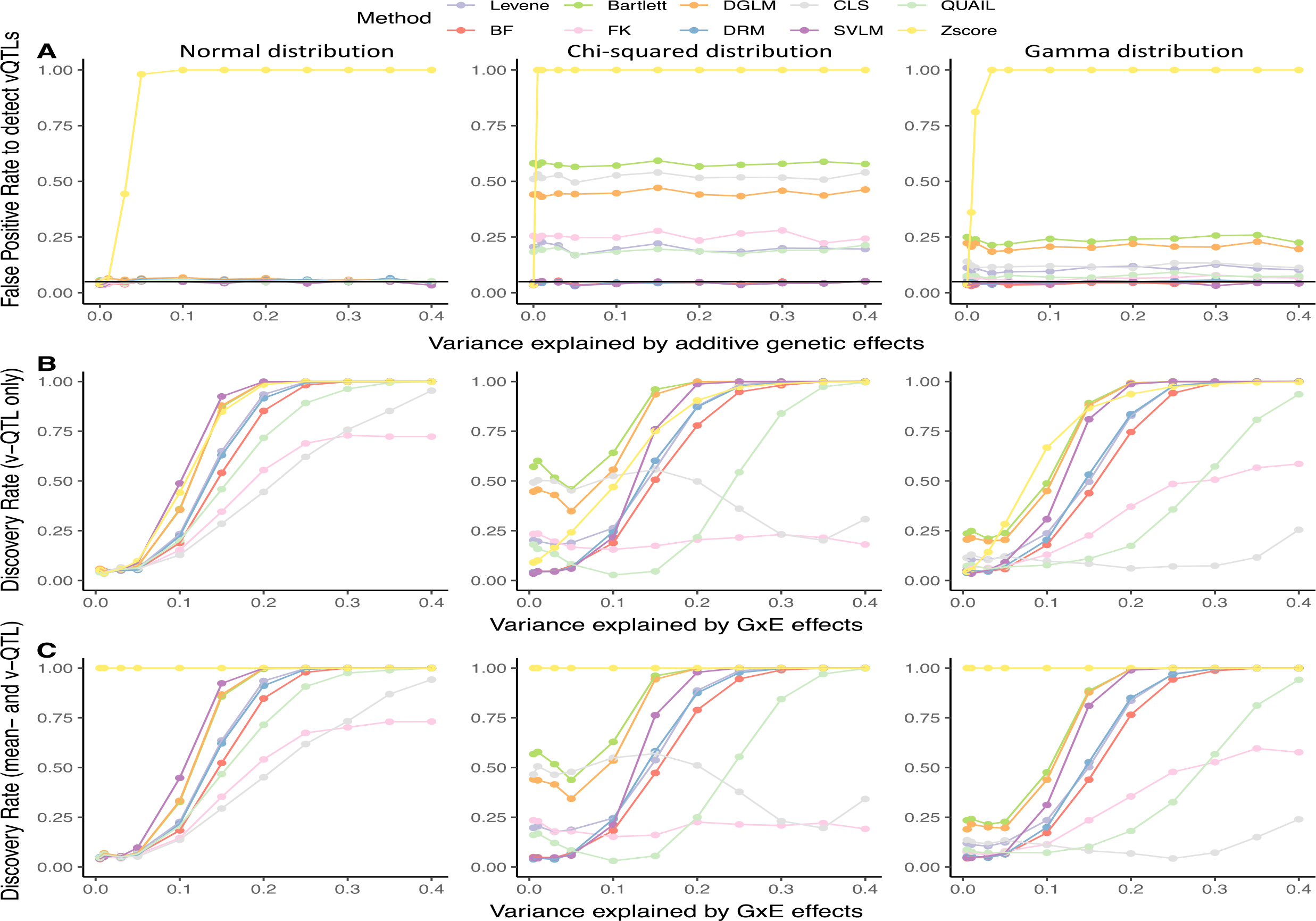
Method performance when the interacting exposure is binary. Discovery rate of methods under the scenario that a single genetic variant affects A) trait level only (*a_mean_* >= 0, *a_var_* = 0), which includes the situation where there is no genetic effect (*a_mean_*= 0, *a_var_* = 0), B) trait variance only (*a_mean_*= 0, *a_var_* > 0), and C) trait level and variance (a*_mean_*= 0.1, *a_var_* > 0). The black horizontal line corresponds to a rate of 0.05. The trait is also affected by noise and environmental factors. The first, second, and third columns represent errors generated from Normal, Chi-squared, and Gamma distributions, respectively.

We observe that the SVLM method performs the best under all trait distributions (Fig 2B, 2C), irrespective of whether there is a mean difference in the phenotypic level across genotype groups. The BF and DRM methods can reach a near 100% discovery rate when the GxE interaction effect size is high, but their discovery rate is lower than that of SVLM. The discovery rates of SVLM, BF, and DRM are higher when the error follows the Normal distribution compared to when the error follows non-Normal distributions. However, the estimated effect size is not accurate for any of the tested methods. The most accurate approach to estimate effect size would be to carry out a follow-up direct test of the GxE interaction (S1 Fig).

In summary, in the presence of a GxE interaction with a discrete exposure, the SVLM, DRM, and BF methods show low FPR, and SVLM performs the best when detecting vQTLs across the ten methods.

### vQTL detection methods comparison for continuous environmental exposures

We next considered GxE interactions that lead to phenotypic variability where the exposure is continuous, such as BMI. We estimated the performance of the 10 methods with continuous exposures which follow the Normal or Uniform distributions. Similar to the methods comparison with discrete exposures, we set the same range of effect sizes for GxE interactions and the errors are again generated from Normal, Chi-squared, or Gamma distributions. The results are obtained for a sample size of 1000 unrelated individuals with a genetic variant at 0.05 MAF. Unlike the discrete exposure scenario where we controlled the sample proportions in the different exposure groups, here we did not control the sample proportions for continuous exposures.

When the exposures follow the Uniform distribution, the Z score method displays the highest FPR, while the SVLM, BF, and DRM methods keep the FPR near to 0.05 (Fig 3A). SVLM outperforms other methods when the error is Normally distributed, but the discovery rate of the SVLM method is lower than that of the BF and DRM methods when the error is non-Normally distributed (Fig 3B). The same findings are observed when the exposure follows the Normal distribution (S2 Fig), which is in contrast to the finding that SVLM is optimal in all data distributions when the exposure is binary (Fig 2B, 2C). The reason could be that both the BF and DRM methods need to calculate the distance between phenotype data values to the phenotype median value per genotype group as input data. When the sample proportions in different exposures are not balanced, many zero values are generated in the input data for these two methods, thus reducing the methods’ ability to model differences among genotype groups. Furthermore, similar to what we observed for binary exposures, the methods’ effect size estimation is not accurate for any of the approaches tested.

**Fig 3.**
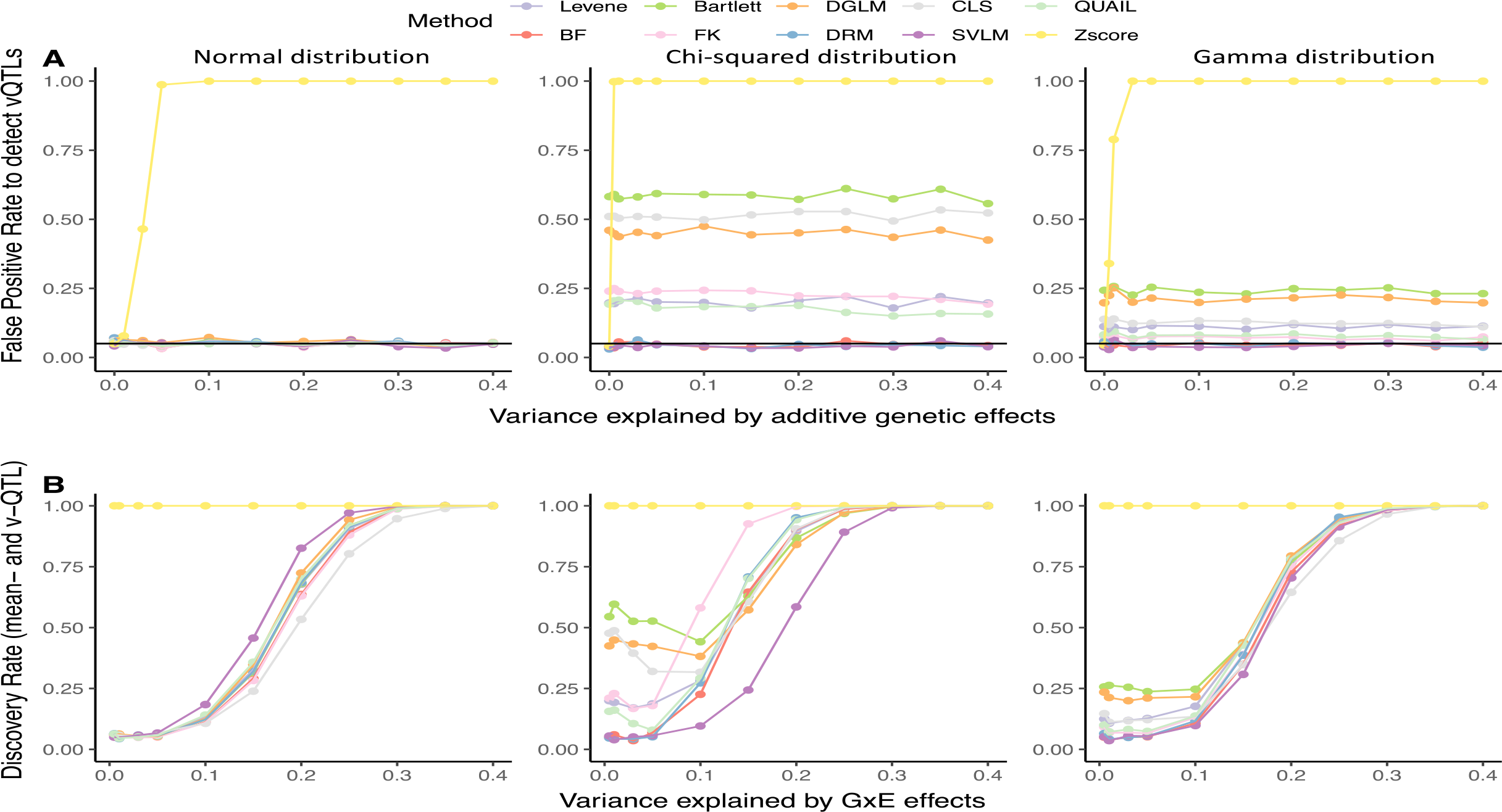
Method performance for Uniformly distributed continuous exposure. Discovery rate of methods under the scenario that a single genetic variant affects A) trait level only (*a_mean_*>= 0, *a_var_* = 0), which includes the situation where there is no genetic effect (*a_mean_* = 0, *a_var_* = 0), and B) trait level and variance (*a_mean_* = 0.1, *a_var_*> 0). The black horizontal line corresponds to a rate of 0.05. The trait is also affected by noise and environmental factors. The first, second, and third columns represent errors generated from Normal, Chi-squared, and Gamma distributions, respectively.

In summary, in the presence of a GxE interaction with a continuous exposure, SVLM, DRM, and BF methods show low FPR. SVLM shows the highest discovery rate when the phenotype data follow the Normal distribution, and DRM performs the best when the phenotype data is non-Normally distributed.

### Phenotype data normalisation may induce mean-variance association across groups

Some vQTL detection methods perform better with Normally distributed phenotypes, for example, Levene’s test, and therefore data normalization may improve method performance. However, data transformations can also induce novel mean-variance associations, where the change of phenotype mean level could lead to a change in the variance of the data distribution [61], for example as observed for the Z score regression model (Fig 2A). To explore this further, we normalized the phenotype data residuals using quantile normalization and repeated the comparison across methods. Taking the binary exposure as an example, all methods showed high FPR for the normalized input data when the error is non-Normally distributed (Fig 4). When the additive genetic effects increase, the FPRs tend to decrease and then increase, forming a U-shape for all methods except SVLM and CLS with non-Normally distributed error. The FPRs after data normalization are greater than the FPRs obtained without data normalization across all methods (Figs 2, 4). Therefore, we do not recommend normalizing phenotype data in vQTL detection.

**Fig 4.**
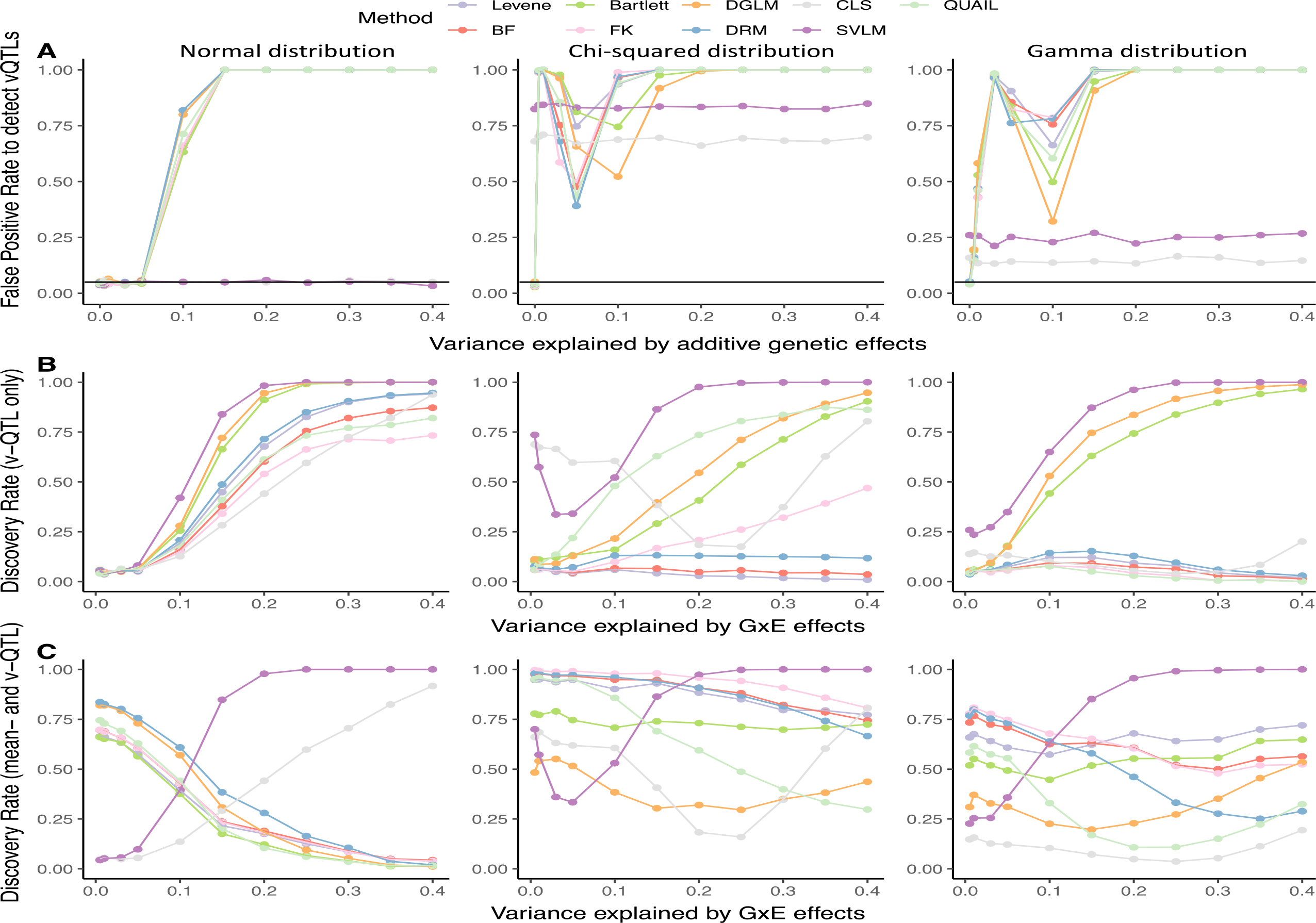
Method performance with rank inverse Normal transformed data. Discovery rates of methods under the scenario that a single genetic variant affects A) trait level only (*a_mean_*>= 0, *a_var_* = 0), which includes the situation where there is no genetic effect (*a_mean_* = 0, *a_var_* = 0), B) trait variance only (*a_mean_* = 0, *a_var_* > 0), and C) trait level and variance (*a_mean_* = 0.1, *a_var_* > 0). The black horizontal line corresponds to a rate of 0.05. The trait is also affected by noise and environmental factors. The first, second, and third columns represent errors generated from Normal, Chi-squared, and Gamma distributions, respectively.

To further interpret this result, we consider that in the simulations under the mean-controlling QTL only scenario (Fig 4A), the trait (Y) is simulated across the three genotype groups as *a_mean_*×0+error (for G_0_), *a_mean_*×1+error (for G_1_), and *a_mean_*×2+error (for G_2_). Therefore, the ranges of Y values are 0 <Y<error (for G_0_), *a_mean_*<Y< *a_mean_*+error (for G_1_), and 2*a_mean_*<Y< 2*a_mean_*+error (for G_2_), respectively. If error>2*a_mean_*or error<*a_mean_*, the variance of the three genotype groups would appear to be different after normalizing the input data Y. Therefore, the mean-controlling QTL will be a spurious signal and the FPR will be high. However, both SVLM and CLS take the genotype as a covariate, so their FPRs do not follow this pattern.

### Power analysis

We estimated power to detect vQTLs under the scenario where the exposure is binary. When a single genetic variant influences trait variability only (*a_mean_* = 0, *a_var_* > 0), the power of DRM and SVLM increases with larger sample sizes and smaller minor allele frequencies. For example, with a MAF of 0.05, a sample size of 1000, and sample proportions in the two exposure categories of 10% and 90%, the power of SVLM and DRM methods could reach 80% when the effect size of the GxE interaction is 0.2 (Fig 5A, S3 Fig). If the sample size increases to 20000, the DRM and SVLM could reach 80% power when the effect size of the vQTL is less than 0.1 (Fig 5A). Furthermore, a more balanced sample distribution across the two exposure categories improves power. Assuming 1000 individuals and equally distributed samples across the two exposure categories, the power of DRM and SVLM surpasses 80% when the effect size of GxE interaction is near 0.1 (Fig 5C).

**Fig 5.**
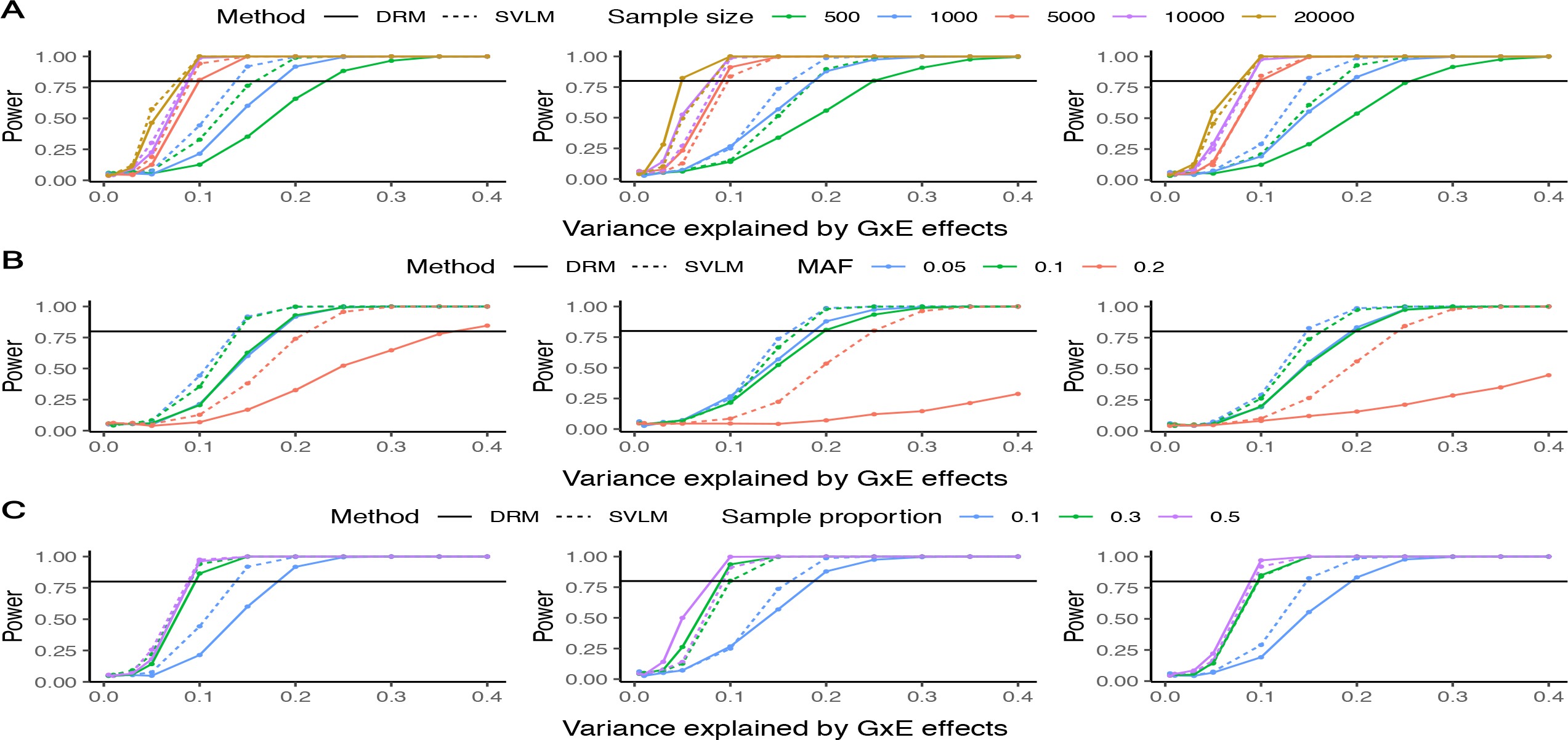
Power analysis when the exposure is binary distributed. Power of BF and DRM across different A) sample sizes, B) MAFs, and C) sample distributions in two exposure categories. The three columns represent situations where the phenotype follows the Normal (left), Chi-squared (middle), and Gamma (right) distributions, respectively.

We also estimated the power of the DRM and SVLM methods when the exposure is continuous. As expected, a larger sample size and smaller MAF again improve the power of DRM and SVLM. When the exposure follows the Uniform distribution, the power of DRM and SVLM reaches nearly 100% with a 20000 sample size, 0.05 MAF, and vQTL effect size of 0.1. When the exposure is Normally distributed, with 20000 samples and a 0.05 MAF, the power of DRM and SVLM could reach at least 80% for a vQTL effect size of 0.25 (S4 Fig).

### Applying DRM and SVLM to human v-eQTL

Two approaches showed the most optimal performance for detecting vQTLs in our simulations, DRM and SVLM. These were next applied to published data to validate previously detected vQTLs. We analysed RNA-seq gene expression profiles from lymphoblastoid cell lines (LCL) from TwinsUK cohort samples to detect variance expression quantitative trait loci (v-eQTLs) identified by two previous studies[32,33]. Briefly, Brown et al. (2014)[33] applied the CLS method to 765 LCL TwinsUK samples where the gene expression data were generated using RNA-seq; and Wang et al. (2014)[32] applied a combination of the FK test and DGLM methods to 825 LCL TwinsUK samples where the gene expression data were generated using Illumina Human HT-12 V3 BeadChips (see Methods). RNA samples partially overlapped across studies, and both reports identified v-eQTLs in these LCL gene expression datasets. Here, we selected 5 genes for which both studies detected v-eQTLs, *NUDT2, BTN3A2, CAPN11, PLXDC2,* and *USP6*, and assessed the performance of DRM and SVLM at these previously reported results.

To this end, we obtained and analysed genotype and LCL RNA-seq gene expression profiles from 765 samples from the TwinsUK cohort. The samples, gene expression data, and imputed genotypes are identical to those in Brown et al (2014)[33]. In addition to DRM and SVLM, we also applied the vQTL detection methods that the original studies used (CLS in Brown et al (2014)[33], and a combination of FK and DGLM in Wang et al. (2014)[32]) to validate the originally published v-eQTL results. Furthermore, both previous studies also reported evidence for epistasis between the identified v-eQTLs and other SNPs (epi-QTLs). Brown et al. (2014)[33] explored the SNPs located within 1Mbp of TSS of the target gene, and Wang et al. (2014)[32] examined genome-wide SNPs. Overall, we selected 5 genes with 10 v-eQTLs and 12 epi-QTLs for validation using DRM, SVLM, CLS, and the combination of FK and DGLM. Multiple testing was taken into account using permutations.

After permutations, we validated 9 of the 10 previously reported v-eQTLs for these 5 genes using both DRM and SVLM (Table 2). The previously reported v-eQTL rs11011705 (v-eQTL for *PLXDC2* in Brown et al. (2014)) did not validate by either method. We also analysed the dataset using the original methods applied by both studies (CLS, FK, DGLM). Altogether, 4 of the 5 v-eQTLs from Brown et al. (2014) were detected at nominal significance using CLS, and 3 surpassed the multiple testing threshold. All 5 of v-eQTLs reported by Wang et al. (2014) validated using DGLM and FK after correcting for multiple testing (FDR<0.05).

**Table2.** Validation of v-eQTLs from Brown et al (2014) and Wang et al (2014). N.S means the statistic of related methods is not significant. ^ represents the method fails to surpass the FDR<0.05.

| Study | Gene | SNP<br>(chr:position) | rsID | Original<br>Sample<br>Size | Validation<br>Sample<br>Size | Validation Methods |  | Original Methods |  |  |
| --- | --- | --- | --- | --- | --- | --- | --- | --- | --- | --- |
|  |  |  |  |  |  | DRM<br>Pvalue | SVLM<br>Pvalue | CLS<br>Pvalue | FK<br>Pvalue | DGLM<br>Pvalue |
| v-eQTLs |  |  |  |  |  |  |  |  |  |  |
| Brown et al | NUDT2 | 9:34318654 | rs10972055 | 765 | 765 | 1.3e-14 | 2.7e-7 | 9.5e-18 | 2.0e-17 | 9.3e-30 |
| Brown et al | BTN3A2 | 6:27024687 | rs12190473 | 765 | 765 | 2.6e-6 | 1.6e-4 | **1.3e-7**<sup>^</sup> | 2.9e-6 | 3.5e-5 |
| Brown et al | CAPN11 | 6:44132782 | rs3757273 | 765 | 765 | 1.6e-11 | 6.3e-10 | 3.1e-24 | 4.3e-11 | 2.7e-29 |
| Brown et al | PLXDC2 | 10:20254725 | rs11011705 | 765 | 765 | **N.S** | **0.04**<sup>^</sup> | **N.S** | **N.S** | 9.4e-8 |
| Brown et al | USP6 | 17:5097573 | rs62072760 | 765 | 765 | 2e-15 | 8.2e-7 | 7.6e-64 | 9.3e-16 | 1.6e-148 |
| Wang et al | NUDT2 | 9:34334015 | rs1337593 | 825 | 765 | 9.6e-15 | 2.7e-7 | 6.7e-18 | 1.3e-17 | 8e-30 |
| Wang et al | BTN3A2 | 6:26415637 | rs742090 | 825 | 765 | 3e-3 | 0.04 | 2e-4 | 1e-4 | 0.03 |
| Wang et al | CAPN11 | 6:44141088 | rs6938938 | 825 | 765 | 6.6e-31 | 4.6e-31 | **3.2e-35**<sup>^</sup> | 1.8e-33 | 8.3e-40 |
| Wang et al | PLXDC2 | 10:20207349 | rs2038912 | 825 | 765 | 3.8e-21 | 6.4e-12 | 6.7e-49 | 6.1e-26 | 7.5e-70 |
| Wang et al | USP6 | 17:5005311 | rs6502843 | 825 | 765 | 2.2e-12 | 1.1e-5 | 3.7e-47 | 1.9e-13 | 5.7e-114 |
| epi-QTLs |  |  |  |  |  |  |  |  |  |  |
| Brown et al | NUDT2 | 9:34381192 | rs3753033 | 765 | 765 | 1.8e-5 | 1.8e-3 | **7.9e-5**<sup>^</sup> | 6.4e-5 | 2e-11 |
| Brown et al | NUDT2 | 9:34256347 | rs10814083 | 765 | 765 | 7e-4 | 9.e-3 | 4.8e-4 | 2e-3 | 3.2e-7 |
| Brown et al | NUDT2 | 9:34284727 | rs7028914 | 765 | 765 | 7.9e-6 | 1.7e-4 | **4.5e-3**<sup>^</sup> | 4.7e-5 | 6.8e-14 |
| Brown et al | BTN3A2 | 6:26276650 | rs9393692 | 765 | 765 | 1e-4 | 5.8e-5 | **6e-3**<sup>^</sup> | **2.1e-3**<sup>^</sup> | 4.1e-9 |
| Brown et al | BTN3A2 | 6:27261324 | rs7746199 | 765 | 765 | 8e-4 | **0.04**<sup>^</sup> | **1.7e-3**<sup>^</sup> | 1.4e-3 | 2.2e-4 |
| Brown et al | CAPN11 | 6:44136899 | rs4711770 | 765 | 765 | 1.6e-5 | 3.2e-4 | 5.9e-8 | 2.8e-4 | 1.6e-17 |
| Brown et al | PLXDC2 | 10:20273583 | rs1111367 | 765 | 765 | 3.5e-13 | 4.6e-6 | 2.5e-31 | 1.5e-18 | 1.2e-25 |
| Brown et al | PLXDC2 | 10:20198883 | rs11594747 | 765 | 765 | **N.S** | **N.S** | **N.S** | **N.S** | **N.S** |
| Brown et al | USP6 | 17:5213391 | rs2585266 | 765 | 765 | 2.2e-33 | 3.8e-19 | 2.4e-50 | 4.6e-47 | Failed to converge |
| Wang et al | NUDT2 | 9:34167694 | rs7032924 | 825 | 765 | **N.S** | **N.S** | **0.01**<sup>^</sup> | **N.S** | **0.02**<sup>^</sup> |
| Wang et al | BTN3A2 | 6:26404374 | rs3799378 | 825 | 765 | 3e-4 | 3.6e-6 | 3.2e-7<sup>^</sup> | 1.6e-4 | 1.4e-5 |
| Wang et al | PLXDC2 | 10:20238864 | rs1326233 | 825 | 765 | 2e-31 | 3.4e-16 | 1.8e-38 | 8.2e-34 | 3.6e-47 |

We next explored the performance of 12 SNPs previously reported by Brown et al. (2014) and Wang et al. (2014) to interact with v-eQTLs, that is, 12 epi-QTLs (Table 2). We identify 10 of these 12 epi-QTLs as v-eQTLs in our dataset by either DRM or SVLM. Two SNPs, rs11594747 (epi-QTL for *PLXDC2* in Brown et al (2014)) and rs7032924 (epi-QTL for *NUDT2* in Wang et al (2014)), are not validated by either method here. However, neither of the 2 epi-QTLs validated when applying the original studies’ methods either (CLS, FK, and DGLM).

Overall, 19 of the 22 tested QTLs (v-QTLs or epi-QTLs) were successfully validated by SVLM or DRM after multiple testing correction. Six of the 22 tested QTLs cannot be validated by CLS, but are validated by at least 3 other methods, which is consistent with our simulation result that CLS shows a lower discovery rate (Fig 2B, 2C). Furthermore, 2 of the 22 tested QTLs were not identified by SVLM, but were detected by DRM, FK, and DGLM (rs732090 and rs7746199). In summary, DRM and SVLM can detect the majority of previously published v-eQTLs and epi-QTLs, and the FK test also shows high replication rates in this dataset.

## Discussion

vQTLs identification could shed light on the genetic basis underlying complex traits. Although multiple approaches have been developed to detect vQTLs, a comprehensive method comparison is lacking. In our study, we used simulated data to evaluate the performance of 10 vQTL detection methods under different biological scenarios of gene-environment interaction. SVLM is optimal when the environmental exposure is discrete. When the environmental exposure is continuous, DRM is optimal in applications to non-Normally distributed data, and SVLM is optimal for Normally distributed phenotypes. A larger sample size, smaller MAF, and more balanced sample distribution in different exposure categories could improve the performance of both SVLM and DRM.

In general, the DRM and SVLM methods display a lower FPR near 0.05 whatever the distributions of data and exposure are. The discovery rate is lower for DRM than for SVLM when the exposure is discrete. We propose that an unbalanced sample distribution across exposure categories affects the performance of DRM in particular because the unbalance would lead to many zero values in the three genotype groups and reduce the power of DRM. Specifically, when a single genetic variant (where G_0_, G_1_, and G_2_ represent three genotype groups) interacts with a binary exposure (E and e), we assume that 90% and 10% of samples are exposed to E and e in each genotype group, respectively. Because most samples are exposed to E, the median phenotype values for the three genotype groups are 0 (90% of values are G_0_×E=0 and 10% of values are G_0_×e=0), E (90% are G_1_×E=E and 10% G_1_×e=e), and 2E (90% are G_2_×E=2E and 10% G_2_×e=2e). Therefore, 90% of the distance between the data point to the median value is 0 in each genotype group, which are the response variables of a linear model in the DRM method. This calculation can be extended to discrete exposures with multiple classes. Further power analysis supports this conclusion that a balanced sample distribution in different categories improves the power of the DRM method. In addition to sample distribution, larger samples size and lower MAF of genetic variants could improve the power of DRM and SVLM as well.

The BF method has often been used to identify vQTL to date (Table 1). Our simulation study showed that the BF method generally shows similar performance to that of DRM, but its discovery rate is slightly lower than DRM under all tested scenarios. However, fewer vQTL studies use SVLM and DRM. The reason might be that to use SVLM one needs to adjust the genotype as a covariate. Therefore, when undertaking genome-wide vWTL detection for one trait, the genotype adjustment will be performed millions of times, which brings a large computational burden. DRM is relatively novel, and so it has not been used in many studies.

The other seven methods, including Levene’s test, Bartlett’s test, FK test, DGLM, CLS, QUAIL, and Z score methods show high FPR when the data follow a non-Normal distribution (Figs 2, 3). The Z score regression method even shows FPR near 1 when the data are Normally distributed. The reason may be that the transformation of the trait to Z scores changes the scale and distribution of original data[61,62]. The data transformation is highly likely to induce mean-variance association, leading to an increased false positive rate.

Our simulation results are consistent with previous comparisons under the assumption of GxE interactions. Wang et al (2019) compared the performance of Bartlett’s, BF, FK, and DGLM methods, and suggested that the BF test outperforms other methods and that any type of data transformation including using squared, cubic, log, or rank inverse normal transformed data would increase the FPR. Marderstein et al (2021) compared 9 methods (DRM, Levene’s test, generalized Levene’s test, BF test, Bartlett’s test, FK test, DGLM, Z score, and SVLM) when sample proportions across discrete exposures are almost balanced. They proposed that the DRM and SVLM show low FPR and DRM is optimal, but they ignored the situation where the sample proportions in different exposure categories are unbalanced. Miao et al (2022)[51] evaluated 4 methods including HLMM, QUAIL, LT, and DRM. They proposed that the HLMM shows inflated FPR, while QUAIL, BF, and DRM methods are good at controlling the FPR. However, our comparison found that QUAIL shows a high FPR. In terms of different simulation settings, Miao et al (2022)’s sample size is larger (200000 versus 1000 in the current study) and they assumed that the trait distribution is less skewed (chi-squared distribution with 6 df *vs* chi-squared distribution with 1 df in the current study). Given these differences, we also evaluated the FPR of QUAIL under the settings where 1/ sample size is 200000 and the trait follows the chi-squared distribution (df=6), which matches Miao et al (2022)[51], 2/ sample size is 200000 and the trait follows chi-squared distribution (df=1), 3/ sample size is 1000 and the trait follows the chi-squared distributions (df=6), and 4) sample size is 1000 and the trait follows the chi-squared distribution (df=1). We found that the FPR of QUAIL is near 0.05 when the chi-squared distribution df equals 6 whatever the sample size is, while the FPR of QUAIL is always higher than those of DRM and SVLM when the df is 1. Therefore, the differences in QUAIL performance are due to differences in the trait distribution, and DRM and SVLM show lower FPR than QUAIL when the trait is largely skewed. Altogether, our results are in line with those obtained from previous studies, and also provide a more detailed comparison under multiple biological scenarios.

In our comparison, we excluded the LRT_MV_, HLMM, BTH, and D test. The original code of LRT_MV_ is missing, HLMM failed to converge when the input data is small, and the BTH and D test require permutation tests. Furthermore, similar to HLMM, the LRT_MV_ calculates the likelihood of genetic effects on the trait level, variance, both or neither to determine how the genetic variant affects the trait distribution. Therefore, like the HLMM, the LRT_MV_ may also fail to converge when the input data is small. Dumitrascu et al. (2019) [58] suggested that the BTH shows similar performance to that of DGLM, and its type I error is lowest in comparison to CLS, Levene’s test, BF, and DGLM tests when the error follows the Gaussian distribution. However, one assumption of the BTH is that the error follows the Gaussian distribution, while its FPR has not been explored for non-Normally distributed traits. The D test compares the accumulated quantile difference of each genotype group. One potential issue is when the trait is largely skewed and the sample size in the three genotype groups is unbalanced. Therefore, the accumulated difference is likely to be different and the FPR might be inflated, although the following permutation tests may help to reduce this inflated error.

We applied the DRM and SVLM methods to validate v-eQTLs from Brown et al (2014)[33] and Wang et al (2014)[32]. The samples, genotype version, and gene expression data were the same as that used in Brown et al (2014)[33], but different from that used in Wang et al (2014)[32]. Altogether, 3 of 5 v-eQTLs and 4 of 9 epi-QTLs detected by Brown et al. (2014)[33] could be validated by CLS in our study, while 4 of 5 v-eQTLs, and 8 of 9 epi-QTLs could be validated by either DRM or SVLM. This is consistent with our result that CLS show a high FPR and low discovery rate. One signal, the previously reported v-eQTL rs11011705 (v-eQTL for *PLXDC2* in Brown et al. (2014)) did not validate by either method. One potential reason for this discrepancy could be the different number of permutations used to control for multiple testing, where Brown et al (2014)[33] used 5 and the current study is based on 20 permutations (for each method, including DRM, SVLM, CLS, FK, and DGLM). The lower number of permutations may introduce spurious signals. All 5 v-eQTLs and 2 of 3 epi-QTL detected by Wang et al (2014) could be validated by FK, DGLM, DRM, and SVLM methods. The FK method performed well in general and was able to validate signals from Brown et al (2014)[33] as well, however, its FPR is near 0.25 when the error is Chi-squared distributed in our simulation. Altogether, DRM and SVLM methods could validate the previously reported v-eQTLs and epi-QTLs.

There are multiple considerations in applications to identify genome-wide vQTLs. One hurdle is how to efficiently conduct vQTL identification across the genome. Currently, none of the proposed methods is computationally efficient, because they either consider the genotype as a covariate (e.g. in SVLM) or calculate the distance of each phenotype data point to a quantile per genotype group (e.g. in DRM), which brings a large computational burden. This is further important because a large sample size is needed to achieve good power for vQTL detection. Given that, the computational time and data storage are challenges in identifying genome-wide vQTLs in large samples.

When interpreting detected signals, it is important to distinguish the true vQTLs from spurious signals. One issue that can impact this is the mean-variance association. The mean level and variance of the data are correlated in data distributions, and this may lead to mean-controlling QTLs being identified as spurious vQTLs. Data transformation is one source of inducing novel mean-variance association because it changes the scale of the phenotype. Therefore, we do not recommend data transformation prior to vQTL detection. Further considerations include linkage disequilibrium (LD) [63–66] between the v-QTL and mean-controlling QTLs. A trait controlled by a mean-controlling QTL (SNP_m_) will display different phenotype levels in each genotype group. If a genetic variant (SNP_1_) is in LD with the SNP_m_, it is possible that the phenotypic variance will be different across genotype groups of SNP_1_ (S5 Fig)[67]. To circumvent this, it would be useful to check LD between known mean-controlling QTLs and detected vQTL. However, it is possible that there are hidden mean-controlling QTLs. Another issue that may induce spurious vQTLs is parent-of-origin or imprinted QTLs. In a parent-of-origin model with a single locus, the phenotypes of the *Aa* genotype category depend on whether the *A* allele is inherited from the father or the mother [68,69]. Therefore, the *Aa* genotype group will display more phenotypic variability than either *AA* or *aa* genotype group in the population. If either homozygote group is missing or has a small sample size, this genetic variant could be detected as a spurious vQTL. Therefore, we suggest setting a minimum threshold for the sample size in the minor homozygote genotype group in real data applications.

In conclusion, we carried out a simulation study to evaluate the performance of ten methods to detect gene-environment interactions as vQTLs. We observed that the DRM and SVLM methods are optimal overall, and can validate the previously detected vQTLs in human gene expression data. We do not recommend data transformation for vQTL detection, because such a transformation may induce a mean-variance association. To improve the power to detect vQTLs, a larger sample size and more balanced sample distribution across exposure categories are helpful. We also suggest follow-ups including LD estimation and comparison with mean-controlling QTLs in the interpretation of vQTLs identified in real data applications.

## Materials and methods

### vQTL mapping approaches

Multiple approaches have been proposed to detect variance QTLs, including variance-only, joint mean-variance, and joint location and scale tests. Here, we consider these approaches for a systematic comparison in a simulation study design.

#### i) Variance-only methods

Tests assessing the heterogeneity of variance are commonly used to detect vQTLs (Fig 1C). The Levene’s test calculates the distance between phenotype data points to the mean phenotypic value of each genotype group and applies a one-way ANOVA test to estimate the variance of distances[43]. A generalized Levene’s scale test was developed without the requirement of independence of samples in Levene’s test[70]. Similarly, the Brown-Forsythe (BF) test calculates the distance from each data point to the median or 10-percent trimmed phenotype mean and then applies a one-way ANOVA to these distances. Compared to the Levene’s test, the BF test was found to be more robust to non-Normally distributed data[44,45]. The Bartlett’s test uses the Chi-squared test to estimate the pooled phenotypic variance of groups, which is one approach to estimating the variance of each group[71]. The Fligner-Killeen (FK) test[47] applies a Chi-squared test to the normalized phenotype rankings of samples, while the prevalent modified FK test compares rankings of the distance between phenotype data points to the group’s median phenotypic value[46]. However, although these methods have been used to detect vQTLs, they also have limitations in that they cannot be used on imputed genotypes and cannot directly account for covariates.

Linear models have also been used to detect vQTLs, and these can take covariates into account (Fig 1C). The squared residual value linear model (SVLM) takes the squared residuals of a linear model, which adjusts the phenotypic trait on covariates and genotype as a response variable [49]. The deviation regression model (DRM) [40] and Z score methods take the distance between phenotype data points to the group’s phenotype median, and Z scores generated from inverse normal transformation as response variables [1,50], respectively. The quantile integral linear model (QUAIL) establishes quantile regression models, because the coefficients of linear models between the genotype and different quantile levels will be different when the genetic variant performs as a vQTL [51]. Brown et al (2014) adjust the phenotype data on genetic effects and take the residuals, and then evaluate the Spearman rank correlation between the squared residuals and genotype (CLS) [33].

#### ii) Mean-variance joint tests

The double generalized linear model (DGLM) iteratively fits two generalized linear mixed models (GLM) which model genetic effects on the phenotypic mean and variance, respectively, until convergence [52,53]. Replacing the GLM with a hierarchical generalized linear model (HGLM) could relax the assumption of the GLM that errors are independent and Normally distributed [54,55]. The LRT_MV_ calculates the maximum likelihood of four linear mixed models, which estimate genetic effects on trait variance, mean, both, and neither, to determine the genetic influence [56]. Likewise, the heteroskedastic linear mixed model (HLMM) calculates the likelihood of genetic effects on trait variance, mean, genetic dominance, all these three effects, and none [57].

Alternative methods include the Bayesian heteroskedastic linear regression model (BTH), which uses the Bayesian framework [58], and the D test which compares the accumulated differences between quantiles of genotype groups (Fig 1C). However, both methods require labor-intensive permutations to determine the appropriate significance threshold, so they are not suitable for genome-wide vQTL mapping [59].

#### iii) Joint location and scale tests

Joint location and scale (JLS) tests combine tests of genetic effects on the mean (for example, linear model) and variance (for example, Levene’s test) to determine whether a genetic variant is associated with a phenotype [48,72,73]. However, JLS results can not specify whether the detected signals act as mean-controlling QTLs or vQTLs. Additional tests are required to interpret the results from JLS [72] in terms of the specific impact on the trait distribution, and therefore JLS tests were not considered in the current simulation design.

In this study we focus on methods that are suitable for genome-wide vQTL mapping without either time-consuming permutations or extra needs to specify whether the identified genetic variants are vQTLs or mean-controlling QTLs. Therefore, we do not consider the above-mentioned BTH, D test, and JLS tests in the comparative simulation analysis. We assume all samples are unrelated, so we also exclude the generalized Levene’s scale test. Furthermore, the original LRT_MV_ code is not available and since its performance was proposed to be similar to that of the DGLM [56], it is excluded from our comparison as well. Lastly, we do not consider HLMM either because we observed that small numerical values can lead to the model failing to converge during testing, and the majority of simulated QTL effects here are small, reflecting published findings. Altogether, this study carries out a simulation-based comparison of 10 vQTL mapping methods, including Levene’s test, BF test, FK test, Bartlett’s test, SVLM, DRM, Z score, CLS, QUAIL, and DGLM.

### Simulation design

We assume that a quantitative phenotype is under the influence of additive genetic effects at a single locus (G), environmental effects from a single exposure (E), GxE interaction effects, and noise (equation 1). We assume that the genetic variant that interacts with environmental factor is the common allele at the single biallelic locus. The exposures E that contributes to the GxE term can be either discrete (e.g. a binary exposure) or continuous (E ∼ Uniform (20, 70) or E ∼ Normal (25,3)). Errors are generated from the Normal, Chi-squared (df=1), and Gamma (shape1=2, shape2=0.5) distributions. We rescaled each term (G, E, GxE, and error) from 0 to 1 with min-max normalization, then we generated the trait Y ensuring that all coefficients (*a_mean_, a_E_, a_var_,* and *a_error_*) sum to 1.

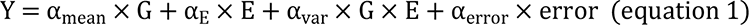

The total sample size (N) and minor allele frequency (MAF) are used to determine the sample size in the three genotype groups. To distinguish vQTLs from genetic variants which are involved in other biological scenarios, such as parent-of-origin effects, we require that there are at least 10 samples in each genotype group. To test different biological scenarios, we set *a_error_* at 0.2. The effect sizes are set at *a_mean_, a_var_* = {0, 0.005, 0.01, 0.03, 0.05, 0.1, 0.15, 0.2, 0.25, 0.3, 0.35, 0.4}. Models are evaluated under situations where the genetic variant impacts, 1/ The phenotype level only (*a_var_* = 0, *a_mean_*≠ 0), 2/ The phenotype variance only (*a_mean_*= 0, *a_var_* ≠ 0), 3/ Both phenotype level and variance (*a_mean_* = 0.1, *a_var_* ≠ 0), or 4/ Neither phenotype level nor variance (*a_var_* = *a_mean_* = 0).

Changes in phenotype variance across groups could lead to differences in the mean phenotype level, which is a hurdle to establishing a variance-only situation. Specifically, if we assume that a trait is influenced by a GxE interaction only, with a single genetic risk variant at a single locus, then there are N_1_, N_2_, and N_3_ unrelated samples in the minor homozygote, heterozygote, and major homozygote group, respectively. Allele G at this locus interacts with exposures which can be classified into k classes (E_1_, E_2_, …, and E_k_). The frequencies of samples that are exposed in each exposure are f_1_, f_2_, …, and f_k_. The mean phenotypic level of the trait in each genotype group is in equation 2.

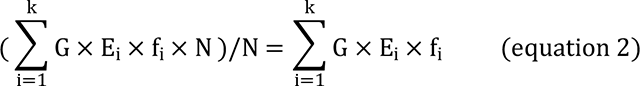

Therefore, the expected mean phenotype levels of the major homozygote, heterozygote, and minor homozygote groups are 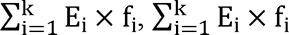, and 0, respectively. The mean phenotype value is influenced by the sample proportions across exposures (*f_i_*) and by the encoded exposures (E_i_). Therefore, a GxE interaction could lead to the shift of the mean of each genotype group. To design a variance only situation, we need to consider sample proportions across exposure categories and how to encode exposures. For example, when the exposure is binary (E and e) and when f_E_ is 10% and f_e_ is 90%, E and e would need to be encoded as 9 and −1 to ensure that there is no difference in the mean phenotype value among the three genotype groups. To simplify our simulation design, we only estimate variance-only situations when the exposure is binary.

We also estimate the power of the methods across a set of parameters, including sample size (N = {500, 1000, 5000, 10000, 20000}) and minor allele frequency (MAF = {0.05, 0.1, 0.2}), and different frequency of samples in the two exposure categories when the exposure is binary (f_E_ = {0.1, 0.3, 0.5}).

### Data pre-processing and method performance assessment

When detecting QTLs, it is often necessary to normalize the phenotype as well as to adjust the phenotype for multiple biological covariates, for example, age, sex, and smoking, as well as technical covariates such as processing batch. Therefore, we adjusted the phenotype for multiple covariates, including the main effect of the hypothesized environmental exposure (E) prior to vQTL detection, as detailed below. For SVLM and CLS, we used a linear model to adjust phenotype levels on both E and genotype (equation 3.1). For the remaining eight methods, we used a linear model to adjust phenotype levels on E (equation 3.2). The resulting residuals and genotype data were then input into each of the ten tested methods to evaluate method performance. To assess the performance of each of the ten tested methods, we estimated the frequency of significant genetic effects (p<=0.05) detected by the method for 1000 simulations. The coefficients of the genotype term in DGLM, SVLM, DRM, QUAIL, and Z score were used as the estimated effect size.

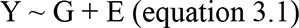

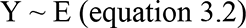

Some methods are also sensitive to the distribution of the phenotype, specifically, Levene’s test, and normalizing the phenotype data (or phenotype residuals after covariate adjustment) may improve model performance[1,42]. Therefore, we also used the normalized residuals as input data for all methods, except for the Z score method which already applies a rank-based inverse normal transformation.

### Model application in real data

We applied a subset of the 10 tested methods to detect previously identified variance QTLs in gene expression data (v-eQTLs). To this end, we used the datasets and results from Brown et al. (2014)[33] and Wang et al. (2014)[32] in detecting v-eQTLs in lymphoblastoid cell line (LCL) samples from twins from the TwinsUK cohort. Brown et al. (2014)[33] used 765 samples and the gene expression profile were generated from RNA-seq, and Wang et al. (2014)[32] used 825 samples and the gene expression data were generated from Illumina Human HT-12 V3 BeadChips. Brown et al. (2014)[33] used the CLS method to detect v-eQTLs after correction for family structure, and they conducted a set of 5 permutation tests for multiple testing corrections, observing 508 associations of v-eQTLs-gene pairs (FDR<0.05). Wang et al. (2014)[32] randomly split twins from the same family into two groups. In one sample group, they applied the FK test and filtered the SNP-probe pairs surpassing P<0.01, and then they used the DGLM method to these filtered SNP-probe pairs to detect v-eQTLs. They conducted 10,000 permutations on each significant SNP-probe pair resulting from DGLM to reduce the false positive rate. Finally, they validated the significant SNP-probe pairs (P_permutation_<0.001) in the other twin sample group. Altogether 99 genes were detected and validated to be associated with at least one cis-v-eQTLs in Wang et al. (2014).

In our study we obtained genotype and gene expression data from LCL samples from 765 samples from the TwinsUK cohort. We obtained the same samples and the same version of genotype imputation and gene expression data as those analysed by Brown et al. (2014)[33]. We focused on 5 genes that were detected to be associated with 5 v-eQTLs in both previous studies, and 5 v-eQTLs and 3 v-eQTLs that were reported to have epistatic effects in Brown et al. (2014)[33] and Wang et al. (2014)[32], respectively. We also considered 9 and 3 genetic variants that were reported to be in epistatic interactions with the v-eQTLs (epi-QTLs) in the original publications, respectively. Altogether, we explored 22 genetic variants and gene expression levels at 5 genes in the current study (Table 2).

The gene expression data were corrected for family structure, zygosity, primer index, age, BMI, GC content, and insert size mode with a linear mixed model. The resulting residuals were then input into CLS and the combination of FK and DGLM to replicate the original findings for v-eQTLs and epi-QTLs. We also applied SVLM and DRM for the detection of v-eQTLs and epi-QTLs. We carried out 20 permutations to calculate the false discovery rate (FDR), and used FDR<0.05 to determine the significance of v-eQTLs and epi-QTLs.

## Acknowledgements

The authors are very grateful to Dr Nick Dand for inputs and critical discussion of the methods and results. XZ acknowledges the financial support of the China Scholarship Council (CSC). This project was supported in part by the JPI ERA-HDHL DIMENSION project (BBSRC BB/S020845/1 to J.T.B.).

## Author contributions

X.Z and J.T.B designed the project. X.Z analysed the project, and the project is supervised by J.T.B. Both authors wrote and approved the final version of the manuscript.

## Code availability

The code for carrying out the simulation study and analysis is available at https://github.com/ZXiaopu/vQTL_sim

## Supporting information

**S1 Fig. Effect size estimation from methods.** When the data follow A) Normal distribution, B) Chi-squared distribution, and C) Gamma distribution.

**S2 Fig. Method performance when the exposure is Normally distributed.** Discovery rates of methods under the scenario that a single genetic variant affects A) trait level only (*a_mean_* >= 0, *a_var_* = 0), which includes the situation where there is no genetic effect (*a_mean_*= 0, *a_var_* = 0), B) trait level and variance (*a_mean_*= 0.1, *a_var_* > 0). The black horizontal line corresponds to a rate of 0.05. The trait is also affected by noise and environmental factors. The first, second, and third columns represent errors generated from Normal, Chi-squared, and Gamma distributions, respectively.

**S3 Fig. Power of methods under different parameter settings when covariates are Uniformly distributed.** A) sample size and B) minor allele frequency. Black horizontal line represents 80% of power. The three columns represent the phenotype data follow Normal (left), Chi-squared (middle), and Gamma (right) distributions, respectively.

**S4 Fig. Power of methods under different parameter settings when covariates are Normally distributed.** A) sample size and B) minor allele frequency. The black horizontal line represents 80% of power.

**S5 Fig. An example of linkage disequilibrium between mean-controlling QTL and vQTL leading to spurious vQTL detection.**

